# Tracking DNA damage localization and chromatin remodeling in live cells using time-resolved quantitative analysis of DNA counterstains

**DOI:** 10.1101/2025.10.01.679728

**Authors:** Greta Paternò, Elisa Longo, Luca Lanzanò

## Abstract

DNA damage profoundly impacts genome stability and cellular homeostasis, and its repair is tightly coordinated with local chromatin remodeling. However, monitoring these rapid chromatin changes in living cells remains challenging. We recently developed QUANDO, an imaging-based method that exploits a simple DNA counterstain to investigate the subnuclear localization of DNA damage in fixed cells. Here, we adapt this approach to track chromatin remodeling at laser-induced DNA damage sites in live cells using Hoechst-based staining. Specifically, PARP1-expressing HeLa cells are exposed to UV laser micro-irradiation in a defined nuclear region, and PARP1 and chromatin dynamics are monitored in real time. We observe that PARP1 rapidly accumulates in the irradiated region but with a heterogeneous pattern: PARP1 initially localizes to high-density chromatin regions (where the concentration of the sensitizer is higher) and gradually redistributes over the whole irradiated region. At the same time, we observe rapid chromatin relaxation, as indicated by decreasing Hoechst intensity and coefficient of variation (CV). In this framework, the PARP inhibitor Talazoparib has the following effects: it slows down PARP1 accumulation, it freezes the PARP1 heterogeneous pattern and blocks chromatin relaxation. Finally, we show that Hoechst-only imaging is sufficient to observe chromatin remodeling: measured chromatin relaxation kinetics are similar in transfected and non-transfected cells, confirming that staining with Hoechst is sufficient for studying chromatin dynamics bypassing the complexities of transfection. These findings underscore the dynamic interplay between DNA damage and chromatin remodeling, demonstrating how conventional nuclear counterstaining can reveal rapid chromatin changes at damage sites, offering new perspectives for investigating genome stability in live cells.

## Introduction

The integrity of the genome is continuously exposed to endogenous and exogenous sources of DNA damage, including reactive oxygen species, replication stress, and environmental agents, which can result in single- and/or double-DNA strand breaks (SSBs and DSBs) (Chatterjee and Walker 2017; Alhmoud et al. 2020). To preserve genomic stability, cells have evolved the DNA Damage Response (DDR), a complex signaling network that detects DNA damage, orchestrates repair pathways, and modulates cell fate decisions. In this process, chromatin remodeling plays a crucial role by facilitating access of repair factors to damaged sites and ensures efficient processing of lesions (Luijsterburg and van Attikum 2011; He et al. 2025).

Among the earliest responses to DNA damage is the activation of poly(ADP-ribose) polymerase 1 (PARP1), which binds DNA breaks and catalyzes the formation of poly(ADP-ribose) chains (PAR) on itself and chromatin-associated proteins such as histones (Izhar et al. 2015; Ray Chaudhuri and Nussenzweig 2017; Smith et al. 2019). PAR signaling critically regulates chromatin architecture: while PARP1 binding may promote local over-compaction, its enzymatic activity induces chromatin relaxation, thereby enhancing DNA accessibility (Sellou et al. 2016; Zong et al. 2022). This dynamic remodeling facilitates the timely recruitment of DNA-binding and repair factors by transiently reshaping the chromatin landscape at sites of damage, ensuring efficient coordination of the repair process (Kruhlak et al. 2006; Smith et al. 2019). Given its pivotal role in DNA repair and chromatin regulation, PARP1 has become a key therapeutic target in tumors with homologous recombination deficiencies, such as BRCA1/2-mutated cancers (Rose et al. 2020), where PARP inhibitors (e.g., Olaparib, Niraparib, Rucaparib, Talazoparib) exert clinical efficacy by trapping PARP on DNA, blocking repair, and promoting the accumulation of cytotoxic lesions (Gunderson and Moore 2015; Póti et al. 2018; Hobbs et al. 2021; Kristeleit et al. 2024).

Over the past decades, several microscopy-based studies have shown that chromatin is a highly dynamic structure undergoing extensive remodeling upon DNA damage, which critically influences repair efficiency and pathway choice (Lakadamyali and Cosma 2015; Lou et al. 2019). To investigate these remodeling events, particularly chromatin change of compaction at damage sites, various imaging strategies have been employed. Among them, a widely used approach involves laser micro-irradiation in combination with a sensitizer to induce localized DNA damage (Zentout et al. 2021). Laser micro-irradiation can be synchronized with time-lapse imaging of live cells to study early events in the DNA damage response. In particular, chromatin remodeling has been observed by monitoring the intensity of photoactivatable histones like H2B-PAGFP (which are photoactivated in the same region of laser-micro-irradiation). In this type of approach, fluorescently labeled chromatin domains expand within seconds to minutes after activation, revealing rapid local relaxation at DNA damage sites (Kruhlak et al. 2006; Burgess et al. 2014; Sellou et al. 2016).

Here, we propose a simpler approach to quantitatively monitor chromatin remodeling using labeling with a vital dye. We demonstrate that DNA damage-induced chromatin reorganization can be effectively studied in live cells using an optimized workflow that combines confocal imaging with a conventional nuclear counterstain. In our approach, we use the DNA counterstain Hoechst 33342 both as a sensitizer and a chromatin remodeling marker. The data are analyzed using a method called QUantitative ANalysis of DNA cOunterstains (QUANDO) (Paternò et al. 2025) that enables the measurement of DNA density level at damage sites. This method combines Image Cross-Correlation Spectroscopy (ICCS) (Comeau et al. 2008; Oneto et al. 2019; Cerutti et al. 2021, 2022) which quantifies the spatial co-distribution between DNA damage markers and chromatin stains, with a DNA density analysis that estimates chromatin compaction levels based on counterstain intensity. Together, ICCS and DNA density offer complementary insights into the local chromatin environment, helping to distinguish between heterochromatin and euchromatin at sites of genomic damage (Oneto et al. 2019; Cerutti et al. 2021, 2022; Paternò et al. 2025). We applied this approach to HeLa cells transfected with a PARP1 chromobody and subjected to UV laser micro-irradiation using Hoechst as a sensitizing dye. First of all, we monitored the dynamics of PARP1 recruitment to ensure that DNA damage was effectively induced: as expected, we observed PARP1 accumulation at damage sites and that this recruitment dynamics was markedly slowed down following treatment with Talazoparib, a potent PARP1 inhibitor. Next, we found that laser-induced DNA damage initially accumulates in high DNA density regions, as indicated by the high values of DNA density and co-localization with heterochromatin that decrease over time. DNA damage is followed by rapid chromatin relaxation, as indicated by a decrease in Hoechst intensity and coefficient of variation (CV). The chromatin relaxation effect is impaired by Talazoparib treatment, which interferes with PARP1-dependent chromatin remodeling. Finally, we found that Hoechst-only imaging provided comparable results in both PARP1-transfected and non-transfected cells, highlighting the method’s robustness and its suitability for single-channel acquisition without requiring transfection.

## Materials and methods

### Cell culture, treatments and labeling

HeLa cells (ATCC n. CCL-2™) are cultured in DMEM (Dulbecco’s modified Eagle’s medium, Gibco™, 11965092) supplemented with 10% Fetal Bovine Serum (Sigma-Aldrich, F9665) and 1% Penicillin/Streptomycin (Sigma-Aldrich, P4333) and maintained at 37°C and 5% CO_2_.

For fluorescence microscopy experiments, 30.000 cells/cm^2^ are seeded on 8-well Ibidi chambered coverslips and incubated at 37°C and 5% CO_2_ for 24 hours to reach approximately 80-90% confluence. Cells are then transiently transfected using Lipofectamine 3000® transfection kit (Thermo Fisher Scientific, L3000-001) according to the manufacturer’s instructions, with a plasmid encoding a PARP1 chromobody tagged with TagRFP (PARP1-RFP) (ChromoTek, xcr).

After transfection, cells are further incubated for 24 hours under the same conditions for subsequent treatments and live-cell imaging. Nuclear staining is performed by incubating the cells with 2 µM Hoechst 33342 (Thermo Fisher Scientific, 62249) for 15 minutes at 37°C, followed by replacement with fresh medium. To inhibit PARP1 activity, cells are treated with 0.5 µM Talazoparib (Selleck Chemicals, BMN 673) for 45 minutes prior to imaging.

### Image acquisition

All measurements are performed on a Leica TCS SP8 confocal laser scanning microscope, using a 63×/1.40 NA oil-immersion objective (HCX PL APO CS2 63/1.40 Oil, Leica Microsystems). For live-cell imaging, the microscope stage was maintained at 37°C with 5% CO_2_ using a temperature- and CO_2_-controlled incubation chamber.

Imaging is carried out using a 512×512 pixel format, line frequency of 8000 Hz (resonant scanner mode), zoom factor of 5.00, a pinhole size set to 1.00 Airy Unit with 10× line accumulation and a pixel size of 72 nm.

Excitation/emission wavelengths are the following: Channel 1 (Ch1) - PARP1-RFP (561/570-650) via hybrid detector operating in photon counting mode (power setting 0.5%); Channel 2 (Ch2) - Hoechst (405/415-510), via hybrid detector operating in photon counting mode (power setting 0.5%).

For the induction of DNA damage, laser micro-irradiation is performed using the 405 nm laser line at 50% power. A rectangular region of interest (ROI) of 18 µm × 4 µm is manually placed at the edge of the nucleus prior to bleaching. The time lapse imaging and bleach setting are the following:

Pre-bleach: 5 iterations, time per iteration = 1.3 sec

Bleach: 2 iterations, time per iteration = 1.3 sec (total exposure time 2.6 sec)

Post-bleach: 50 iterations, time per iteration = 3 sec.

In the time-lapse analysis, we define the beginning of the micro-irradiation as the time t = 0 so that t =2.6 sec corresponds to the first image acquired post-irradiation.

### Data processing

The acquired images are pre-processed on Fiji (Schindelin et al. 2012) to obtain the suitable input files for DNA density analysis and Image Cross-Correlation Spectroscopy (ICCS), with the QUANDO algorithm implemented in MATLAB (https://github.com/llanzano/QUANDO), as thoroughly described in Paternò et al. 2025. The input files are the following:

- “Nuclei selection count masks”: label images identifying each nucleus as a distinct object over time in the time-lapse sequence, used to define nuclear boundaries. They are generated as follows: the images of the DNA channel (Ch2) from the time-lapse sequence of the same nucleus are converted into binary images using the function ‘‘Threshold’’ of ImageJ, using the “Default” threshold algorithm. The nuclei are identified and listed as objects using the ‘‘Analyze particles’’ function and the images of the ‘‘Count Masks’’ are saved;
- “DNA damage area count masks”: label images representing the intersection between nuclei and the micro-irradiated ROI, representing the irradiated nuclear areas;
- “Binary masks of DNA damage foci”: binary images highlighting pixels corresponding to occurrence of DNA damage. The binary masks of DNA damage are generated by excluding background pixels from the PARP1 channel (Ch1), setting a fixed intensity threshold of 80% of the maximum intensity value;
- “Intensity images”: background-subtracted fluorescence images, used for quantitative intensity measurements. The intensity images are generated by smoothing the images and subtracting the background from the intensity images of nuclei channel (Ch2) using the function “Subtraction of Background” (rolling ball radius of 100 pixels). The rolling ball radius size corresponds approximately to the maximum size of heterochromatin features that we observed in our images.

### Image Cross-Correlation Spectroscopy (ICCS) analysis

The Image Cross-Correlation Spectroscopy (ICCS) analysis is based on a modified version of the ICCS algorithm (Oneto et al. 2019) (https://github.com/llanzano/ICCS), well described in (Cerutti et al. 2022; Privitera et al. 2024). The algorithm is performed in MATLAB (The MathWorks, Natick, Massachusetts). We adapt the algorithm to perform automatic calculation of ICCS parameters on each time point of the time-lapse. For each time point, the main algorithm output is the parameter f_1_ (f_2_) values which represent the fraction of signal in channel 1 (channel 2) which is cross-correlated with the other channel. Values of this parameter range from 1 (maximum cross-correlation), to 0 (no cross-correlation), to −1 (maximum anti-correlation) (Cerutti et al. 2022).

The colocalization fraction extracted by ICCS is analogous but not identical to the Pearson Correlation Coefficient (Cerutti et al. 2022). The main difference is that the ICCS parameter is calculated using the image cross- and auto-correlation functions. Specifically, the cross-correlation fraction f_1_ is defined as the ratio between the amplitude (estimated at zero spatial lag) of the cross-correlation function and the amplitude (extrapolated at zero spatial lag) of auto-correlation function of channel 2.

In this application, we cross-correlate the signal corresponding to DNA damage (PARP1 channel) with the signal corresponding to the DNA marker (Hoechst channel). Thus, the value of the parameter f_1_ represents the fraction of colocalization of DNA damage with regions of high DNA density (heterochromatic regions). The analysis is limited to the irradiated area of the nucleus using “DNA damage area count masks” as the region of analysis.

### DNA density analysis

For each time point, we calculate the image of the normalized DNA intensity as:

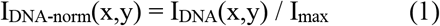

Where I_DNA_(x,y) is the intensity image of the DNA channel and I_max_ is the maximum value of I_DNA_(x,y) in the “Nuclei selection count masks”.

Then we calculate the average value of the normalized DNA intensity in the region of interest (ROI):

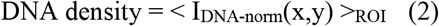

Where the ROI is represented by the intersection between the “Nuclei selection count masks” and the region defined by “Binary masks of DNA damage foci”. For each time-point, the value of “DNA density” represents the average value of the normalized DNA intensity, calculated at the DNA damage foci. We expect high values of DNA density if DNA damage overlaps with heterochromatic regions.

### Intensity and CV analysis

For each time-point, average values of PARP1 intensity (Ch1) and Hoechst intensity (Ch2) are calculated in the ROI defined by the “DNA damage area count masks”. The coefficient of variation (CV) of the intensity in the DNA channel is calculated as (Martin et al. 2021) :

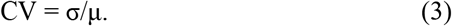

Where σ and μ are the standard deviation and the mean value of Hoechst intensity in the same ROI, respectively.

### Graphs and statistical analysis

Graphs and statistical analyses are performed using GraphPad Prism version 8.0.0 for Windows, GraphPad Software, San Diego, California USA, www.graphpad.com. For all the analyses, differences among two groups are analyzed by T-test. The values are expressed as mean ± s.d. in Fig. 2-5, mean ± s.e.m. in Fig. 3-4, and a p < 0.05 is accepted as significant.

For the intensity time trace analysis, the accumulation of PARP1-RFP (T_on_) at damage sites is examined by fitting the short-term intensity profiles (average intensity within the DNA damage region) using the exponential one-phase association model in GraphPad Prism. The intensity profile during the decay phase (T_off_) is analyzed separately using the exponential one-phase decay equation.

## Results

### Hoechst-based approach to investigate chromatin dynamics following DNA damage

To investigate chromatin remodeling at DNA damage sites in living cells, we established an experimental workflow combining Hoechst nuclear staining, laser micro-irradiation, time-lapse imaging and a quantitative image analysis (**Fig. 1**). Notably, Hoechst also acts as a photosensitizer: upon UV excitation, it generates reactive oxygen species, enabling the induction of site-specific DNA damage in live cells (Hurst and Gasser 2019). As illustrated schematically in **Fig. 1a**, the method can be applied to PARP1-transfected and non-transfected cells. In both cases, nuclei were labeled with Hoechst, and localized DNA damage was induced by a focused 405 nm laser within a defined nuclear region (dashed box). Time-lapse imaging allowed simultaneous monitoring of PARP1 recruitment to damage sites (only in PARP1-transfected cells) and chromatin remodeling over time (**Fig. 1b**). Regions of interest (ROI) were defined around the irradiated area to track the evolution of fluorescence signals.

**Figure 1.**
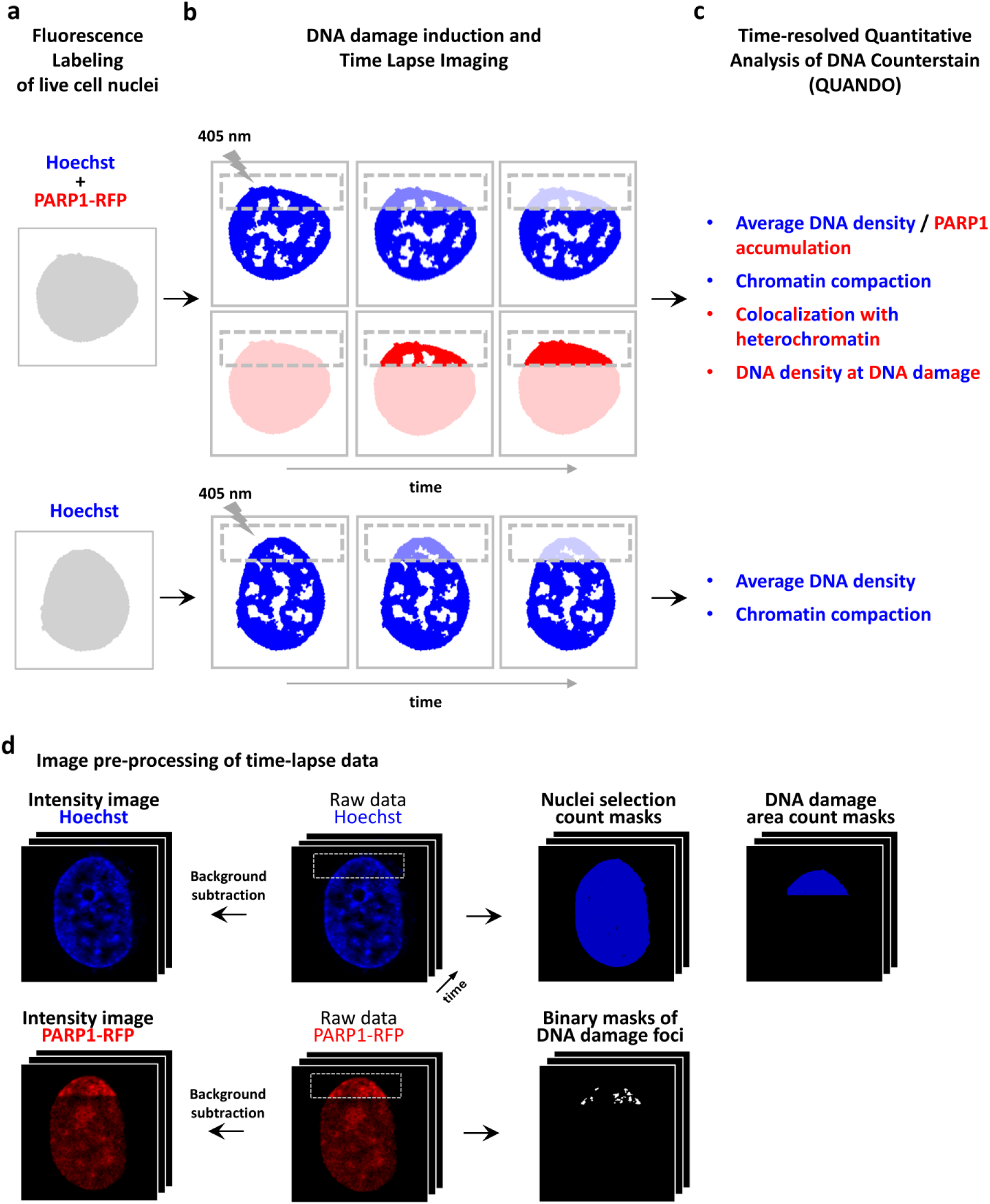
Schematic overview of the experimental workflow used to monitor chromatin remodeling at DNA damage sites in live cells. (a) Live cells were stained with Hoechst (blue) and included both PARP1-transfected (red) and non-transfected cells. (b) DNA damage was induced with 405 nm laser micro-irradiation in a specific nuclear region (dashed box), and time-lapse imaging was used to monitor PARP1 recruitment and chromatin dynamics. (c) Acquired images were processed with QUANDO to quantify DNA damage localization (ICCS and DNA density analysis) and chromatin remodeling (Hoechst intensity and CV). (d) Schematic description of the processing of time-lapse raw data and preparation of the images required to perform the analysis.

Following image acquisition, we processed the data using a time-resolved QUANDO analysis. In PARP1-transfected cells we evaluated DNA damage localization by: (i) measuring the colocalization of DNA damage with heterochromatin regions using Image Cross-Correlation Spectroscopy (ICCS), and (ii) analyzing changes in local chromatin density through normalized Hoechst intensity at the damaged area as shown in (Paternò et al. 2025). We expect higher colocalization and DNA density values when DNA damage occurs in heterochromatic regions, and lower values when damage localizes to euchromatin. In parallel, we quantified the average PARP1 signal intensity as a readout of PARP1 recruitment, providing insights into the protein’s dynamic response to DNA damage, including under conditions where PARP activity is inhibited by Talazoparib (see below). In both transfected and non-transfected cells, we evaluated chromatin remodeling, using the Hoechst signal intensity and its coefficient of variation (CV). The CV reflects the heterogeneity of nuclear staining: higher Hoechst intensity and CV values are typically associated with more compact chromatin, while lower values indicate chromatin relaxation (**Fig. 1c**). The whole time-resolved analysis is automated thanks to the generation of count masks that identify the region of analysis for each time point (**Fig. 1d**).

### PARP1 accumulation following laser-induced DNA damage

In order to use Hoechst both as a sensitizer and a chromatin marker, we set up a configuration in which the laser micro-irradiation procedure did not cause significant photobleaching of the Hoechst dye. To this end, we compared Hoechst fluorescence intensity immediately before and 2.6 sec after micro-irradiation. No significant changes were detected, as confirmed qualitatively by confocal images (**Fig. 2a**) and quantitatively by normalized intensity plots (**Fig. 2b**). Nevertheless, laser irradiation effectively induced DNA damage, as demonstrated by PARP1 accumulation at the irradiated region of interest (ROI) (**Fig. 2c**). To capture a broad chromatin landscape and better visualize spatial organization, the ROI was designed with a rectangular shape (approx. 72 µm^2^), encompassing an extended nuclear area. Before the micro-irradiation (pre-irradiation), confocal imaging revealed that in control cells (**Fig. 2c**), PARP1 was uniformly distributed throughout the nucleoplasm, with a higher concentration observed in the nucleoli, in keeping with previous reports (Buchfellner et al. 2016; Longo et al. 2024). Immediately after damage induction (2.6 sec post-irradiation), PARP1 rapidly accumulated at the targeted site, forming distinct high-intensity foci. By the end of the time-lapse acquisition (149.6 sec post-irradiation), the fluorescence signal appeared more diffuse and evenly distributed across the ROI. In contrast, in cells pre-treated with the PARP inhibitor Talazoparib (**Fig. 2e**), PARP1 dynamics was altered. Although the initial accumulation at the damage site resembled that of the control cells, by the end of the time-lapse acquisition the fluorescence signal remained confined to well-defined foci, preserving a structured pattern. This behavior suggests a delayed yet more stable binding of PARP1 in the presence of the inhibitor.

**Figure 2.**
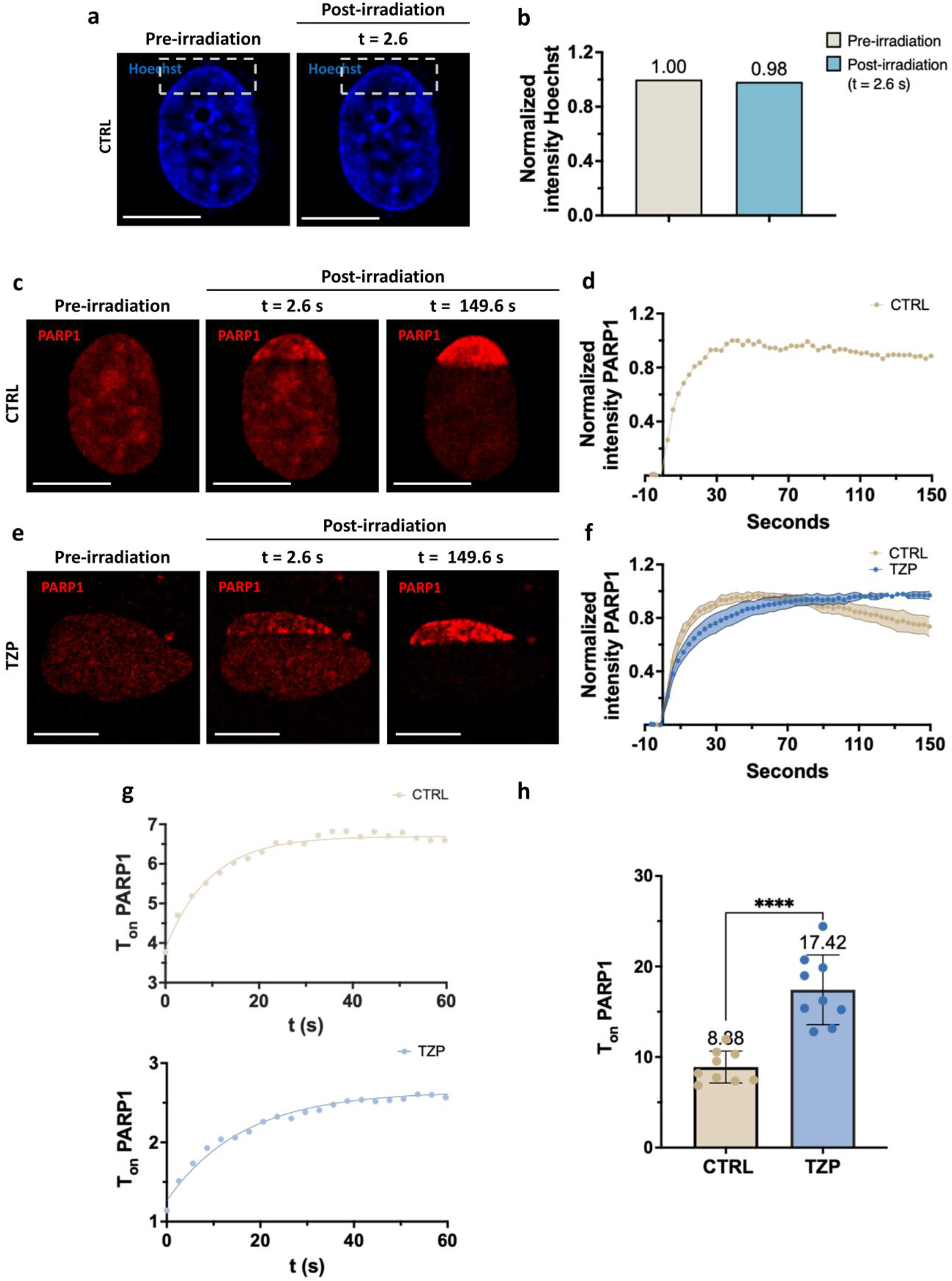
Time-lapse imaging of PARP1 dynamics following DNA damage. (a) Representative images of live HeLa cell nuclei stained with Hoechst before and immediately after irradiation (t = 2.6 sec). Dashed box represents the defined nuclear region targeted by laser-induced DNA damage. Scale bars: 10 µm. (b) Comparison of normalized intensity Hoechst between pre-irradiation and post-irradiation. T-Test was applied and no significant difference were detected. (c–e) Confocal images show PARP1 recruitment at the damage site in control (CTRL) and Talazoparib-treated cells (TZP). For each condition, three time points are shown: pre-irradiation, and post-irradiation at t = 2.6 sec and t = 149.6 sec. Scale bar: 10 µm. (d–f) Normalized intensity profiles of PARP1 over time in control and Talazoparib-treated cells. (f) Cloud plots represent mean ± s.d. from 9 cells. (g) Representative fitting curves using a one-phase association model (T_on_) of PARP1 accumulation in control and Talazoparib-treated cells within the first 60 sec. h) Dot plots showing the comparison of PARP1 T_on_ values between control and Talazoparib-treated cells. Data points represent mean ± s.d. from 9 cells, T-Test ****p < 0.0001.

To characterize these dynamics, fluorescence intensity values of PARP1 were calculated over time. In **Fig. 2d**, we reported the normalized intensity profile of PARP1 in a single untreated control cell. **Fig. 2f** showed the averaged intensity profiles of untreated (beige) and Talazoparib-treated (light blue) cells. In control cells, PARP1 intensity rapidly increased, reached its peak at 53.6 sec, the normalization reference point, and then gradually declined. In contrast, Talazoparib-treated cells displayed a delayed peak at 134.6 sec, with a slower accumulation rate and no subsequent decrease, resulting in a sustained plateau. These observations indicated that PARP1 binding was delayed but exhibited increased stability in the presence of the inhibitor.

To quantitatively assess the recruitment kinetics, we fitted the intensity curves using a single-exponential function (**Fig. 2g**), which estimates the characteristic time required for PARP1 accumulation at DNA damage sites. The recruitment time constant (T_on_) was significantly higher in Talazoparib-treated cells (light blue), with T_on_ = 17.42 sec ± 3.84 (mean ± s.d. from 9 cells), compared to untreated controls (beige), which showed a T_on_ = 8.88 sec ± 1.76 (mean ± s.d. from 9 cells), as reported in **Fig. 2h**. The slower yet sustained accumulation is consistent with the PARP trapping effect, in which Talazoparib impairs PARP1 dissociation from DNA damage sites and promotes its prolonged retention on chromatin, thereby slowing down the accumulation kinetics.

### Time-resolved QUANDO-based quantification of PARP1 localization after laser-induced DNA damage

Next, we applied the QUANDO method to analyze the localization of laser-induced DNA damage in control and Talazoparib-treated cells. Specifically, we tracked the recruitment of endogenous PARP1 over time following 405 nm laser micro-irradiation. PARP1 localization was evaluated using the parameters: (i) DNA density, and (ii) colocalization with heterochromatin (see Methods). As previously demonstrated by (Paternò et al. 2025), high DNA density values combined with strong colocalization with heterochromatin indicate that DNA damage preferentially localizes within more compact chromatin regions. Conversely, low DNA density and reduced colocalization suggest a redistribution of the damage signal into euchromatic regions.

As shown in **Fig. 3a-b**, confocal images acquired immediately after laser micro-irradiation (t = 2.6 sec) revealed that PARP1 (red) rapidly accumulated at the damage site and preferentially localized to regions of high Hoechst intensity (blue), which corresponded to more compact chromatin. Then, control cells displayed a progressive diffusion of the PARP1 signal, which became less confined and more evenly distributed across the damaged area, suggesting that, as the DNA damage response progressed, PARP1 spread into lower-density, euchromatic regions. In contrast, Talazoparib-treated cells maintained a more restricted and structured PARP1 distribution, indicating sustained accumulation within dense chromatin compartments.

**Figure 3.**
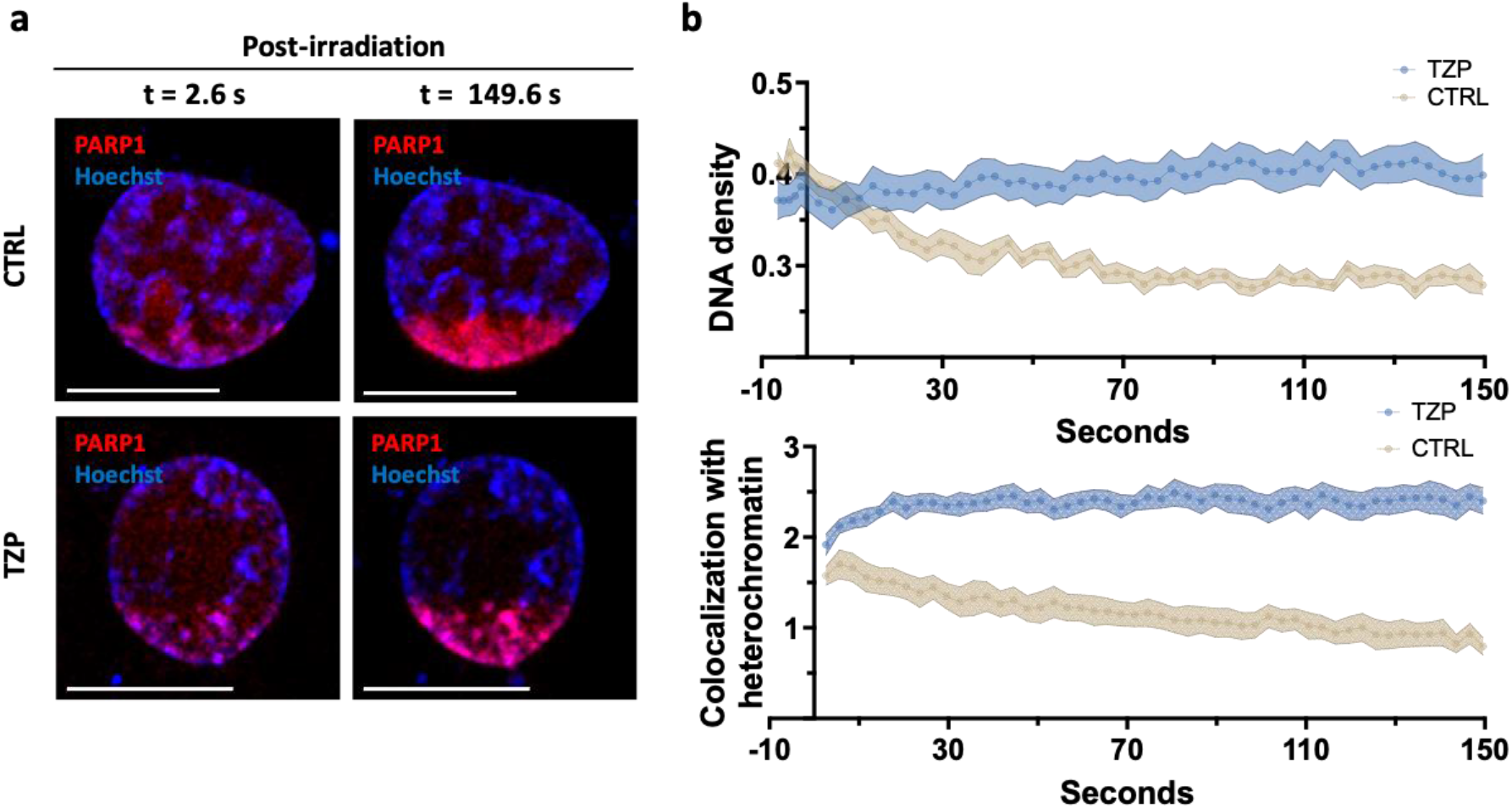
PARP1 localization dynamics revealed by time-resolved QUANDO. (a) Representative confocal images of control (CTRL) and Talazoparib-treated (TZP) HeLa cells at early (t = 2.6 sec) and late (t = 149.6 sec) time points after laser-induced DNA damage. Images show PARP1 (red) and DNA counterstaining with Hoechst (blue). Scale bars: 10 µm. (b) Top: DNA density corresponding to PARP1 accumulation over time in control (beige) and Talazoparib-treated (blue) cells. Cloud plots show the mean ± s.e.m. of 8 cells. Bottom: Colocalization of PARP1 with heterochromatin over time in control (beige) and Talazoparib-treated (blue) cells. Pattern cloud plots show the mean ± s.e.m. of 8 cells.

Within the first ∼10 seconds, in both control (beige) and Talazoparib-treated cells (light blue) PARP1 is preferentially localized in high-density chromatin regions, confirming its initial recruitment to heterochromatin, likely due to the elevated local concentration of the Hoechst sensitizer (higher DNA density and higher concentration of AT-rich sequences) that concentrates the occurrence of DNA damage in these regions. As time progressed, control cells displayed a marked redistribution of PARP1 toward lower-density regions, along with a decrease in colocalization with heterochromatin, consistent with its dynamic relocation into euchromatin. In contrast, Talazoparib-treated cells maintained PARP1 enrichment in high-density regions with sustained colocalization with heterochromatin, showing minimal redistribution over time. This suggests that PARP inhibition traps PARP1 at the sites of damage within heterochromatin and impairs the progression of downstream repair mechanisms. Overall, these results highlight how chromatin context influences PARP1 dynamics and how PARP inhibitors interfere with its chromatin remodeling-dependent redistribution.

### Hoechst-based confocal analysis captures chromatin remodeling dynamics in live cells

We next analyzed the Hoechst channel alone in both PARP1-transfected cells (**Fig. 4a-b**) and non-transfected cells (**Fig. 4c-d**). In PARP1-transfected cells (**Fig. 4a**), confocal imaging revealed a clear chromatin relaxation at the site of laser-induced DNA damage, visible as a localized reduction in Hoechst intensity that became progressively more homogeneous starting from the earliest post-irradiation time point (t = 2.6 sec) until the final time point (t = 149.6 sec). This relaxation pattern was absent in Talazoparib-treated cells, where chromatin remained visibly compacted in the damaged region and displayed a distinct pattern of euchromatic and heterochromatic subdomains.

**Figure 4.**
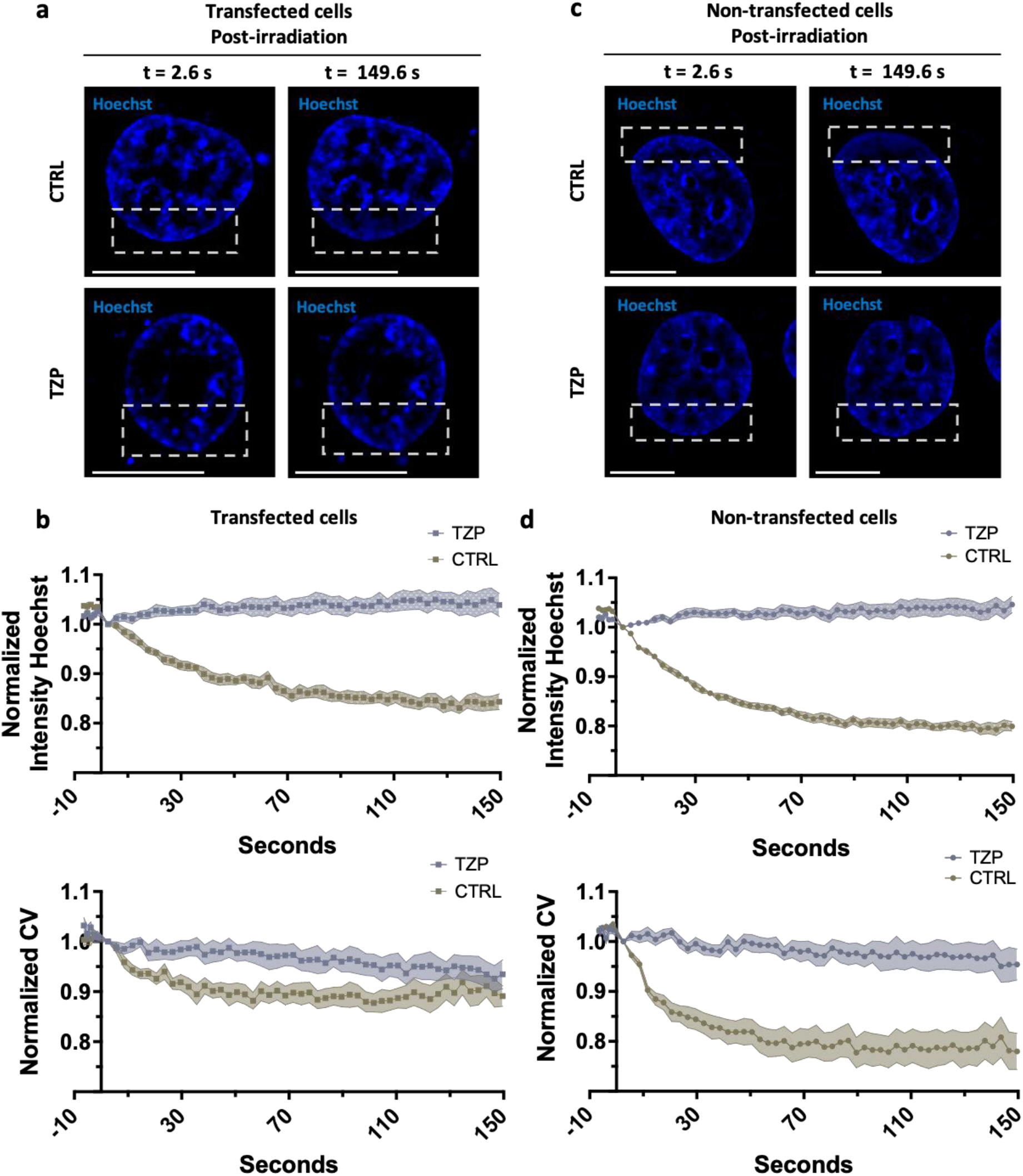
Single-channel imaging reveals chromatin relaxation after DNA damage and its inhibition by Talazoparib in both PARP1-transfected and non-transfected cells. (a) Representative confocal images of PARP1-transfected HeLa cells at post-irradiation time points (t = 2.6 sec and 149.6 sec) following laser-induced DNA damage (dashed square). Conditions include control (CTRL) and Talazoparib-treated cells (TZP). Staining: Hoechst (blue) and PARP1 (not shown). Scale bars: 10 µm. (b) PARP1-transfected cells. Top: Quantitative analysis of normalized intensity of Hoechst in control (beige square) and Talazoparib-treated (gray square) cells. Pattern cloud plots show the mean ± s.e.m. of 8 cells. Bottom: Normalized coefficient of variation (CV) in control (beige square) and Talazoparib-treated (gray square) cells. Cloud plots show the mean ± s.e.m. of 8 cells. (c) Representative confocal images of non-transfected cells at post-irradiation time points (t = 2.6 sec and 149.6 sec) following laser-induced DNA damage (dashed square). Conditions include control and Talazoparib-treated cells. Staining: Hoechst (blue). Scale bars: 10 µm. (d) Non-transfected cells. Top: Quantitative analysis of normalized intensity of Hoechst in control (beige circle) and Talazoparib-treated (gray circle) cells. Pattern cloud plots show the mean ± s.e.m. of 8 cells. Bottom: Normalized CV in control (beige circle) and Talazoparib-treated (gray circle) cells. Cloud plots show the mean ± s.e.m. of 8 cells.

Chromatin reorganization within the damaged region was quantitatively assessed using Hoechst-based metrics (**Fig. 4b**). Specifically, we monitored both the average Hoechst intensity and its coefficient of variation (CV), which together provide a readout of chromatin compaction. Higher values of these metrics correspond to more compact chromatin, whereas lower values indicate chromatin relaxation. As shown in Fig. 4b, values were normalized to the first post-irradiation time point. In control cells (beige squares), both Hoechst intensity and CV progressively decreased over time in the DNA damage region, consistent with dynamic chromatin relaxation. This trend was altered in Talazoparib-treated cells (grey squares), where Hoechst intensity remained stable and the CV showed only a minor reduction, suggesting that PARP inhibition restrains chromatin decompaction in response to DNA damage.

A similar response was observed in non-transfected cells (**Fig. 4c–d**). Quantitative analysis of Hoechst intensity and CV further confirmed these observations: in control cells (beige circle), both parameters decreased progressively, whereas in Talazoparib-treated cells (grey circle), they remained largely unchanged. All together, these observations demonstrate that even single-channel analysis of nuclear DNA staining provides informative and quantifiable insight into chromatin dynamics, offering a simple and minimally invasive readout to monitor structural changes in live cells. Notably, comparable results were obtained in both PARP1-transfected and non-transfected cells, underscoring the robustness and general applicability of this approach.

### Chromatin relaxation kinetics are comparable in transfected and non-transfected cells

Finally, we quantified the decay kinetics of Hoechst intensity and CV over time in both control PARP1-transfected (**Fig. 5a**) and non-transfected cells (**Fig. 5b**). Specifically, we calculated the T_off_ values (exponential decay constants) to quantify the kinetics of chromatin relaxation and to directly compare the temporal dynamics between PARP1-transfected and non-transfected cells. As shown in **Fig. 5a**, the average T_off_ for Hoechst intensity was 40.7 sec in PARP1-transfected cells (grey) and 34.7 sec in non-transfected cells (light blue), as illustrated by the representative fits. The difference was not statistically significant, indicating that the kinetics of Hoechst signal decay, reflecting chromatin decompaction, were comparable between the two conditions. Similarly, in **Fig. 5b**, the T_off_ values for the coefficient of variation were 16.9 sec in PARP1-transfected cells (violet) and 25.4 sec in non-transfected cells (blue). These comparable values suggest that the dynamics of chromatin heterogeneity within the damaged region evolve at a similar pace in both cell populations. Importantly, this validation confirms that informative measurements can be obtained using the Hoechst channel alone, without the need for PARP1-RFP transfection, thereby simplifying the workflow and extending the applicability of this approach. These data also confirm that expression of the PARP1 chromobody (Buchfellner et al. 2016) does not interfere with the chromatin relaxation process that follows DNA damage induction.

**Figure 5.**
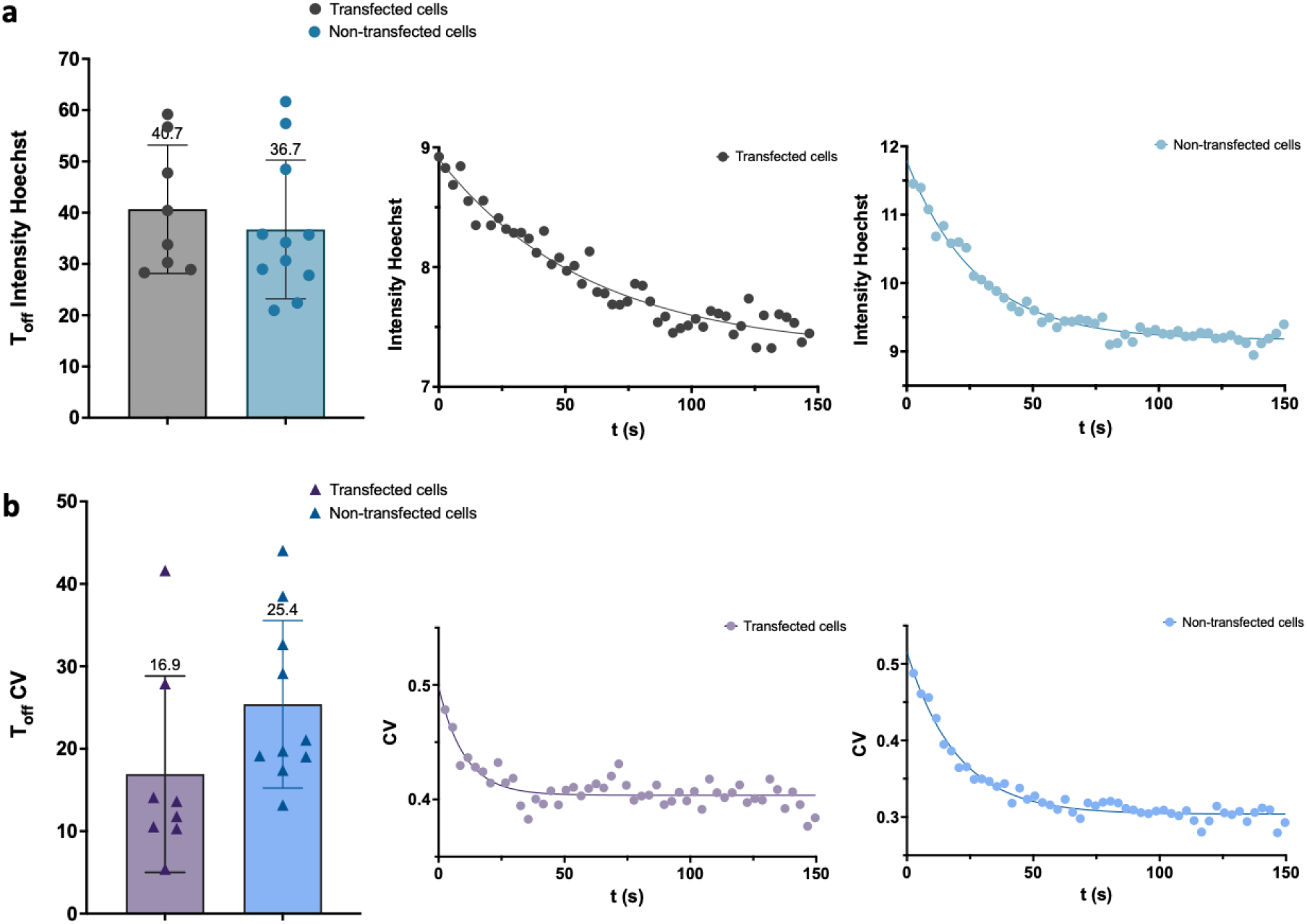
Comparable chromatin relaxation dynamics in PARP1-transfected and non-transfected cells. (a) Left: Dot plots showing the comparison of decay constants (T_off_) for Hoechst intensity in PARP1-transfected (grey circle) and non-transfected (light blue circle) cells. Data points represent the mean ± s.d of 8 PARP1-transfected cells and 10 non-transfected cells. T-Test was applied and no significant difference were detected. Right: Representative decay fitting curves of Hoechst intensity over time in PARP1-transfected (grey circle) and non-transfected (light blue circle) cells. (b) Left: Dot plots showing the comparison of decay constants (T_off_) for Hoechst CV in PARP1-transfected (violet triangle) and non-transfected (blue triangle) cells. Data points represent the mean ± s.d of 8 PARP1-transfected cells and 10 non-transfected cells. T-Test was applied and no significant difference were detected. Right: Representative decay fitting curves of Hoechst CV over time in PARP1-transfected (violet circle) and non-transfected (blue circle) cells.

## Discussion/Conclusions

DNA damage elicits a well-characterized cascade of chromatin remodeling events that promote lesion accessibility, coordinate the recruitment of repair factors, and ensure genome stability (Burgess et al. 2014). These early chromatin changes, most notably the relaxation of chromatin at the damage site, are a hallmark of the DNA damage response (DDR). However, it remains unclear whether, and to what extent, such remodeling events occur uniformly (Shaban and Seeber 2020).

To study these dynamics in live cells, we set up a relatively simple approach based on confocal microscopy and staining with a Hoechst dye used both as a sensitizer for laser micro-irradiation and a marker of chromatin remodeling. By carefully optimizing dye concentration and illumination parameters, we defined a protocol to induce DNA damage with minimal photobleaching of the Hoechst dye, enabling reliable live-cell monitoring of chromatin remodeling via changes in Hoechst fluorescence intensity and coefficient of variation (CV).

To investigate the pattern of laser-induced DNA damage, we applied a time-resolved version of our QUANDO method, recently published by Paternò et al. 2025. QUANDO previously showed that in U937-PR9 cells, spontaneous and PML-RARα–induced damage localizes mainly to euchromatin, while Neocarzinostatin-induced damage is more evenly distributed (Paternò et al. 2025). Here, we found that PARP1 initially accumulated in more compact chromatin domains and later redistributed to euchromatic regions in control cells, while remaining confined to dense chromatin regions under Talazoparib. Control cells showed progressive chromatin relaxation at damage sites, indicated by decreased Hoechst intensity and coefficient of variation (CV), whereas Talazoparib blocked this relaxation. Similar chromatin dynamics in PARP1-transfected and non-transfected cells confirmed that Hoechst staining alone effectively monitors chromatin remodeling. Overall, chromatin remodeling occurs dynamically after DNA damage but is hindered by PARP inhibition, and Hoechst analysis offers a minimally invasive live-cell method to track these changes.

Previous studies have investigated chromatin remodeling during DNA damage response using various microscopy-based approaches. For instance, Izhar et al. 2015 reported, similarly to our observations, that UV laser micro-irradiation combined with Hoechst staining induces localized chromatin relaxation (called “antistripes” by the authors) mediated by PARP1. However, their analysis was qualitative and did not provide spatial and temporal quantification (Izhar et al. 2015). Several studies have exploited labeling of histones with photoconvertible fluorescent proteins and time-lapse imaging (Kruhlak et al. 2006; Burgess et al. 2014; Sellou et al. 2016). Fluorescence correlation spectroscopy (FCS) has been used to quantify the mobility of inert or chromatin-associated proteins within the chromatin environment (Hinde et al. 2014; Lou et al. 2020; Longo et al. 2025). Förster Resonance Energy Transfer (FRET) imaging has been used to measure chromatin compaction at the nanoscale following induction of DNA damage (Lou et al. 2019; Pelicci et al. 2020). In this framework, our method provides a simpler, yet quantitative, way to analyze chromatin remodeling in live cells during the early DNA damage response.

Looking ahead, this approach could be combined with live-cell super-resolution imaging techniques, such as image scanning microscopy (ISM) or Airyscan microscopy, to enhance spatial resolution while preserving live-cell compatibility and minimizing phototoxicity (Korobchevskaya et al. 2017; Castello et al. 2019; Di Franco et al. 2025). Its versatility also makes it suitable for application in more physiologically relevant systems, including 3D spheroids that better recapitulate tissue architecture and nuclear organization. Overall, this work opens new avenues to dissect the interplay between chromatin architecture and DNA repair dynamics with minimal experimental perturbation.

## Statements & Declarations

### Author contributions

GP and LL designed the study and wrote the manuscript. GP and EL prepared samples and collected data. LL wrote the software. GP and LL analyzed data and discussed results. All authors critically reviewed the manuscript.

## Data availability

All datasets are provided within the main text and can be obtained from the corresponding authors upon request. This study includes no data deposited in external repositories.

## References

Alhmoud JF, Woolley JF, Al Moustafa A-E, Malki MI (2020) DNA Damage/Repair Management in Cancers. Cancers (Basel) 12:1050. 10.3390/cancers12041050

Buchfellner A, Yurlova L, Nüske S, et al (2016) A New Nanobody-Based Biosensor to Study Endogenous PARP1 In Vitro and in Live Human Cells. PLoS One 11:e0151041. 10.1371/journal.pone.0151041

Burgess RC, Burman B, Kruhlak MJ, Misteli T (2014) Activation of DNA Damage Response Signaling by Condensed Chromatin. Cell Rep 9:1703–1717. 10.1016/j.celrep.2014.10.060

Castello M, Tortarolo G, Buttafava M, et al (2019) A robust and versatile platform for image scanning microscopy enabling super-resolution FLIM. Nat Methods 16:175–178. 10.1038/s41592-018-0291-9

Cerutti E, D’Amico M, Cainero I, et al (2021) Evaluation of sted super-resolution image quality by image correlation spectroscopy (QuICS). Sci Rep 11:20782. 10.1038/s41598-021-00301-x

Cerutti E, D’Amico M, Cainero I, et al (2022) Alterations induced by the PML-RARα oncogene revealed by image cross correlation spectroscopy. Biophys J 121:4358–4367. 10.1016/j.bpj.2022.10.003

Chatterjee N, Walker GC (2017) Mechanisms of DNA damage, repair, and mutagenesis. Environ Mol Mutagen 58:235–263. 10.1002/em.22087

Comeau JWD, Kolin DL, Wiseman PW (2008) Accurate measurements of protein interactions in cells via improved spatial image cross-correlation spectroscopy. Mol Biosyst 4:672. 10.1039/b719826d

Di Franco E, Tedeschi G, Scipioni L, et al (2025) Exploiting the detector distance information in image scanning microscopy by phasor-based SPLIT-ISM. Biomed Opt Express 16:1270. 10.1364/BOE.551255

Gunderson CC, Moore KN (2015) Olaparib: an oral PARP-1 and PARP-2 inhibitor with promising activity in ovarian cancer. Future Oncology 11:747–757. 10.2217/fon.14.313

He L, Moon J, Cai C, et al (2025) The interplay between chromatin remodeling and DNA double-strand break repair: Implications for cancer biology and therapeutics. DNA Repair (Amst) 146:103811. 10.1016/j.dnarep.2025.103811

Hinde E, Kong X, Yokomori K, Gratton E (2014) Chromatin Dynamics during DNA Repair Revealed by Pair Correlation Analysis of Molecular Flow in the Nucleus. Biophys J 107:55–65. 10.1016/j.bpj.2014.05.027

Hobbs EA, Litton JK, Yap TA (2021) Development of the PARP inhibitor talazoparib for the treatment of advanced BRCA1 and BRCA2 mutated breast cancer. Expert Opin Pharmacother 22:1825–1837. 10.1080/14656566.2021.1952181

Hurst V, Gasser SM (2019) The study of protein recruitment to laser-induced DNA lesions can be distorted by photoconversion of the DNA binding dye Hoechst. F1000Res 8:104. 10.12688/f1000research.17865.2

Izhar L, Adamson B, Ciccia A, et al (2015) A Systematic Analysis of Factors Localized to Damaged Chromatin Reveals PARP-Dependent Recruitment of Transcription Factors. Cell Rep 11:1486– 1500. 10.1016/j.celrep.2015.04.053

Korobchevskaya K, Lagerholm B, Colin-York H, Fritzsche M (2017) Exploring the Potential of Airyscan Microscopy for Live Cell Imaging. Photonics 4:41. 10.3390/photonics4030041

Kristeleit R, Leary A, Oaknin A, et al (2024) PARP inhibition with rucaparib alone followed by combination with atezolizumab: Phase Ib COUPLET clinical study in advanced gynaecological and triple-negative breast cancers. Br J Cancer 131:820–831. 10.1038/s41416-024-02776-7

Kruhlak MJ, Celeste A, Dellaire G, et al (2006) Changes in chromatin structure and mobility in living cells at sites of DNA double-strand breaks. J Cell Biol 172:823–834. 10.1083/jcb.200510015

Lakadamyali M, Cosma MP (2015) Advanced microscopy methods for visualizing chromatin structure. FEBS Lett 589:3023–3030. 10.1016/j.febslet.2015.04.012

Longo E, Paternò G, Diaspro A, Lanzanò L (2025) Measuring PARP1 mobility at DNA damage sites by segmented fluorescence correlation spectroscopy. Biophys J. 10.1016/j.bpj.2025.05.013

Longo E, Scalisi S, Lanzanò L (2024) Segmented fluorescence correlation spectroscopy (FCS) on a commercial laser scanning microscope. Sci Rep 14:17555. 10.1038/s41598-024-68317-7

Lou J, Priest DG, Solano A, et al (2020) Spatiotemporal dynamics of 53BP1 dimer recruitment to a DNA double strand break. Nat Commun 11:5776. 10.1038/s41467-020-19504-3

Lou J, Scipioni L, Wright BK, et al (2019) Phasor histone FLIM-FRET microscopy quantifies spatiotemporal rearrangement of chromatin architecture during the DNA damage response. Proceedings of the National Academy of Sciences 116:7323–7332. 10.1073/pnas.1814965116

Luijsterburg MS, van Attikum H (2011) Chromatin and the DNA damage response: The cancer connection. Mol Oncol 5:349–367. 10.1016/j.molonc.2011.06.001

Martin L, Vicario C, Castells-García Á, et al (2021) A protocol to quantify chromatin compaction with confocal and super-resolution microscopy in cultured cells. STAR Protoc 2:100865. 10.1016/j.xpro.2021.100865

Oneto M, Scipioni L, Sarmento MJ, et al (2019) Nanoscale Distribution of Nuclear Sites by Super-Resolved Image Cross-Correlation Spectroscopy. Biophys J 117:2054–2065. 10.1016/j.bpj.2019.10.036

Paternò G, Scalisi S, Dellino GI, et al (2025) Location of oncogene-induced DNA damage sites revealed by quantitative analysis of a DNA counterstain. European Biophysics Journal. 10.1007/s00249-025-01755-x

Pelicci S, Tortarolo G, Vicidomini G, et al (2020) Improving SPLIT-STED super-resolution imaging with tunable depletion and excitation power. J Phys D Appl Phys 53:234003. 10.1088/1361-6463/ab7cf8

Póti Á, Berta K, Xiao Y, et al (2018) Long-term treatment with the PARP inhibitor niraparib does not increase the mutation load in cell line models and tumour xenografts. Br J Cancer 119:1392–1400. 10.1038/s41416-018-0312-6

Privitera AP, Scalisi S, Paternò G, et al (2024) Super-resolved analysis of colocalization between replication and transcription along the cell cycle in a model of oncogene activation. Commun Biol 7:1260. 10.1038/s42003-024-06972-2

Ray Chaudhuri A, Nussenzweig A (2017) The multifaceted roles of PARP1 in DNA repair and chromatin remodelling. Nat Rev Mol Cell Biol 18:610–621. 10.1038/nrm.2017.53

Rose M, Burgess JT, O’Byrne K, et al (2020) PARP Inhibitors: Clinical Relevance, Mechanisms of Action and Tumor Resistance. Front Cell Dev Biol 8:. 10.3389/fcell.2020.564601

Schindelin J, Arganda-Carreras I, Frise E, et al (2012) Fiji: an open-source platform for biological-image analysis. Nat Methods 9:676–682. 10.1038/nmeth.2019

Sellou H, Lebeaupin T, Chapuis C, et al (2016) The poly(ADP-ribose)-dependent chromatin remodeler Alc1 induces local chromatin relaxation upon DNA damage. Mol Biol Cell 27:3791–3799. 10.1091/mbc.E16-05-0269

Shaban HA, Seeber A (2020) Monitoring global chromatin dynamics in response to DNA damage. Mutation Research/Fundamental and Molecular Mechanisms of Mutagenesis 821:111707. 10.1016/j.mrfmmm.2020.111707

Smith R, Lebeaupin T, Juhász S, et al (2019) Poly(ADP-ribose)-dependent chromatin unfolding facilitates the association of DNA-binding proteins with DNA at sites of damage. Nucleic Acids Res 47:11250–11267. 10.1093/nar/gkz820

Zentout S, Smith R, Jacquier M, Huet S (2021) New Methodologies to Study DNA Repair Processes in Space and Time Within Living Cells. Front Cell Dev Biol 9:. 10.3389/fcell.2021.730998

Zong W, Gong Y, Sun W, et al (2022) PARP1: Liaison of Chromatin Remodeling and Transcription. Cancers (Basel) 14:4162. 10.3390/cancers14174162

